# A long-acting GDF15 analog causes robust, sustained weight loss and reduction of food intake in an obese non-human primate model

**DOI:** 10.1101/2022.12.05.519153

**Authors:** Songmao Zheng, David Polidori, Yuanping Wang, Brian Geist, Xiefan Lin-Schmidt, Jennifer L. Furman, Serena Nelson, Andrea R. Nawrocki, Simon A. Hinke

**Author notes:** Author contributions: S.A.H., S.Z., and A.R.N. designed research; S.A.H., S.Z., D.P., J.L.F., S.N., Y.W., B.G. and X. L-S. performed research; S.A.H., S.Z., D.P., Y.W. and B.G. analyzed data; S.A.H., S.Z., D.P., and A.R.N. wrote the paper. Submitting Author: Simon Hinke, Ph.D., Janssen R&D, 1400 McKean Rd, Spring House, PA 19477.

## Abstract

GDF15 is a circulating polypeptide associated with cellular stress, and recently linked to metabolic adaptation. GDF15 has a half-life of approximately 3 hours in and acts at the GFRAL receptor selectively expressed in the area postrema. To characterize the effects of sustained GFRAL agonism on food intake (FI) and body weight (BW), we developed a half-life extended analog of GDF15 (Compound H; CpdH) suitable for reduced dosing frequency and tested its effects in obese cynomolgus monkeys. Animals were treated once weekly for 12 weeks with 0.048, 0.16, or 1.6 mg/Kg of CpdH or with 0.02 mg/Kg of the long-acting GLP-1 analog dulaglutide as a positive control. FI was measured daily and BW was measured biweekly. Mechanism-based longitudinal exposure-response (E-R) modeling was performed to characterize the effects of CpdH and dulaglutide on FI and BW. The integrated novel model accounts for both acute, exposure-dependent effects of treatments to reduce FI and the compensatory changes in energy expenditure (EE) and FI that occur over time in response to weight loss. CpdH had approximately linear, dose-proportional pharmacokinetics with a half-life of ≈8 days and treatment with CpdH led to dose- and exposure-dependent reductions in FI and BW. The 1.6 mg/Kg CpdH dose reduced mean FI by 57.5% at 1 week and provided sustained FI reductions of 31.5% from weeks 9-12, leading to a peak reduction in BW of 16±5%. Dulaglutide had more modest effects on FI (reductions between 15-40%) and peak BW loss was 3.8±4.0%. Longitudinal modeling of both the FI and BW profiles suggested reductions in BW observed with both CpdH and dulaglutide were fully explained by the exposure-dependent reductions in FI without any increase in EE. Upon verification of pharmacokinetic/pharmacodynamic relationship established in monkeys and human for dulaglutide, we predicted that CpdH could reach double digit BW loss in human. In summary, treatment with a long-acting GDF15 analog led to sustained dose- and exposure-dependent reductions in food intake in a monkey model of obesity and holds potential for effective clinical obesity pharmacotherapy.

**Significance Statement:** GDF15 activation of GFRAL receptors in the hindbrain controls food intake and body weight. Here we describe the effect and durability of a circulating half-life extended analog of GDF15 (Compound H) on food intake and body weight loss in a spontaneously obese cynomolgus monkey model. Inclusion of a translational treatment arm with a weight loss agent, dulaglutide, permitted pharmacokinetic/pharmacodynamic modeling and comparison of both GDF15 and GLP-1 based weight loss mechanisms, and development of an allometric scaling based mathematical model to estimate the efficacy of Compound H in human obese subjects.

## Introduction

Obesity is a public health epidemic afflicting approximately 39% of the global population, with disproportionately greater impact in the Americas and Europe (1). Body Mass Index greater than 25 Kg/m^2^ is associated with increased risk of heart disease, stroke, type 2 diabetes (T2DM) and cancer – leading causes of premature preventable death (2). As the prevalence of obesity grows, so does the associated health care burden, recently estimated to be $85.7 billion dollars each year (3). Although evidence is accumulating that significant weight loss may reduce development of comorbidities (4), and evaluation of cardiovascular impacts of existing weight loss and diabetes therapies have shown promise, few pharmacological interventions have been approved for the treatment of obesity. Limited efficacy, poor tolerability and insurance coverage further reduce broader use of approved agents (4). Hence, considerable efforts are underway to elucidate novel and complementary weight loss mechanisms for clinical development.

Growth Differentiation Factor-15 (GDF15) is a secreted circulating polypeptide associated with energy balance (5, 6). Whereas animals deficient in GDF15 are prone to increased body weight and fat mass (7), elevated GDF15 levels in rodents derived from xenograft or transgene overexpression induce a lean phenotype with improved metabolism, resisting diet-induced obesity and insulin resistance (8–10). Secreted GDF15 is released from peripheral tissues in response to cellular stress and tissue injury and acts centrally in the area postrema (AP). Intracerebroventricular injection of GDF15 in mice decreases food intake (FI), and lesion of these hindbrain regions abolishes the effect of exogenous GDF15 on body weight (BW) and diet consumption (11). Additional evidence for a central mechanism of GDF15 was suggested by c-Fos activation in AP and hypothalamic neurons following peripheral injection of the hormone (11). Indication of relevance of this biological function of the GDF15 mechanism in humans was first proposed from the correlation of elevated GDF15 and reduced body mass index in patients with advanced cancers; similarly, a negative correlation between BW and circulating GDF15 was reported in monozygotic twins (12, 13). More recently, reduced FI and concomitant BW reduction in response to long-acting GDF15 analogs have been described in spontaneously overweight cynomolgus monkeys (14, 15), indicating this mechanism is conserved up the phylogenetic tree.

Recently, GFRAL (GDNF Family Receptor Alpha-Like), a distant member of the Glial-derived Neurotrophic Factor (GDNF) receptor family (16), was independently de-orphaned by several drug discovery laboratories as the cognate receptor for GDF15. Ligand binding by heterologous expression of GFRAL was shown to be selective for GDF15, with members of the TGFβ superfamily and GDNF sub-family unable to bind GFRAL or compete with GDF15 binding (14, 17–19). Co-receptor RET tyrosine kinase association was required for GDF15 intracellular signaling through GFRAL, however, it was not required for receptor binding (14, 18, 19). Reports confirmed expression of GFRAL in the AP in mouse, rat, non-human primate and human (14, 17–19), consistent with earlier studies implicating these brain regions in GDF15’s activity (11). Generation of global GFRAL knockout mice confirmed that GFRAL receptor expression is required for GDF15 to exert effects on food intake, body weight, and glucose tolerance in obese mice fed a high fat diet (14, 17–19). At the same time, it was confirmed that lack of GFRAL did not alter the effect of GLP-1 receptor agonism on food intake, body weight or glucose tolerance (14, 17).

Given the conservation of the GDF15/GFRAL mechanism in non-human primates (14, 15), in the current study, we sought to evaluate the durability of the effects of a long acting GDF15 agonist, Compound H (CpdH), on food intake and body weight in an obese monkey model. In addition, we sought to develop a mechanistic model characterizing the longitudinal and exposure-dependent effects of treatment on food intake and body weight to facilitate predictions of effective doses for future human clinical evaluation. As a translational control, we included dulaglutide, a long-acting glucagon-like peptide-1 (GLP-1) agonist, to help assess the translatability of the spontaneously obese NHP animal model and the modeling-based predictions of efficacious exposures using a clinically approved drug with well-characterized weight loss in humans. A mechanism-based pharmacokinetic/pharmacodynamic (PK/PD) model was derived using mean PK, energy intake (EI, calculated based on total calorie data from documented food consumption; used interchangeably as food intake/FI) and BW data, and permitted simulation of effects in humans based upon publicly available data for dulaglutide. The model has broad applicability for allometric scaling weight loss agents from NHP models to anticipated human clinical effects, enabling setting clinical endpoints and aiding in trial design.

## Results

### Compound Serum Exposure

The cohort of spontaneously obese cynomolgus monkeys used in the current study had an average baseline BW of 9.1±0.2 Kg, BMI of 46.0±1.0 Kg/m^2^, and 15.1±1.5% total body fat (n=32; **Supplementary Table 1**). Grouped baseline characteristics of the cohort are found in **Supplementary Table 2**.

On day 0, animals were injected subcutaneously with vehicle (8% Sucrose, 0.04% polysorbate20, pH6.5), CpdH (0.048, 0.16, or 1.6 mg/Kg, selected based on PK modeling to represent estimates of food intake EC_10_, EC_30-50_ and EC_90_ once steady state exposures were achieved) or 0.02 mg/Kg dulaglutide (dose selected to approximate average human clinical exposure); PK parameters are shown in **Table 1**. Dose-dependent increases in serum CpdH were observed, achieving steady-state exposure within 2-3 weeks (**Figure 1A**). Dulaglutide exposure was maintained within the human clinical range, but with a higher peak to trough ratio (**Figure 1B**). Whereas no animals demonstrated loss of exposure to dulaglutide throughout the study, several CpdH treated animals showed rapid loss of serum exposure resulting in exclusion from the study from that point forward, as per the data analysis plan (see **Methods**). Rapid loss of compound exposure is consistent with induction of immune-mediated clearance, likely due to species differences in HSA or GDF-15 sequences. All animals in the low dose group lost detectable CpdH by day 51 of treatment; serum exposure of CpdH was lost in 4 animals in the 0.16mg/Kg dose group by day 70 and 3 animals in the high dose CpdH group lost exposure during the first month of treatment (**Figure 1C**). Animal number in **Figure 1C** represents the N of each data point in **Figures 2** and **3** over time.

**Table 1:**
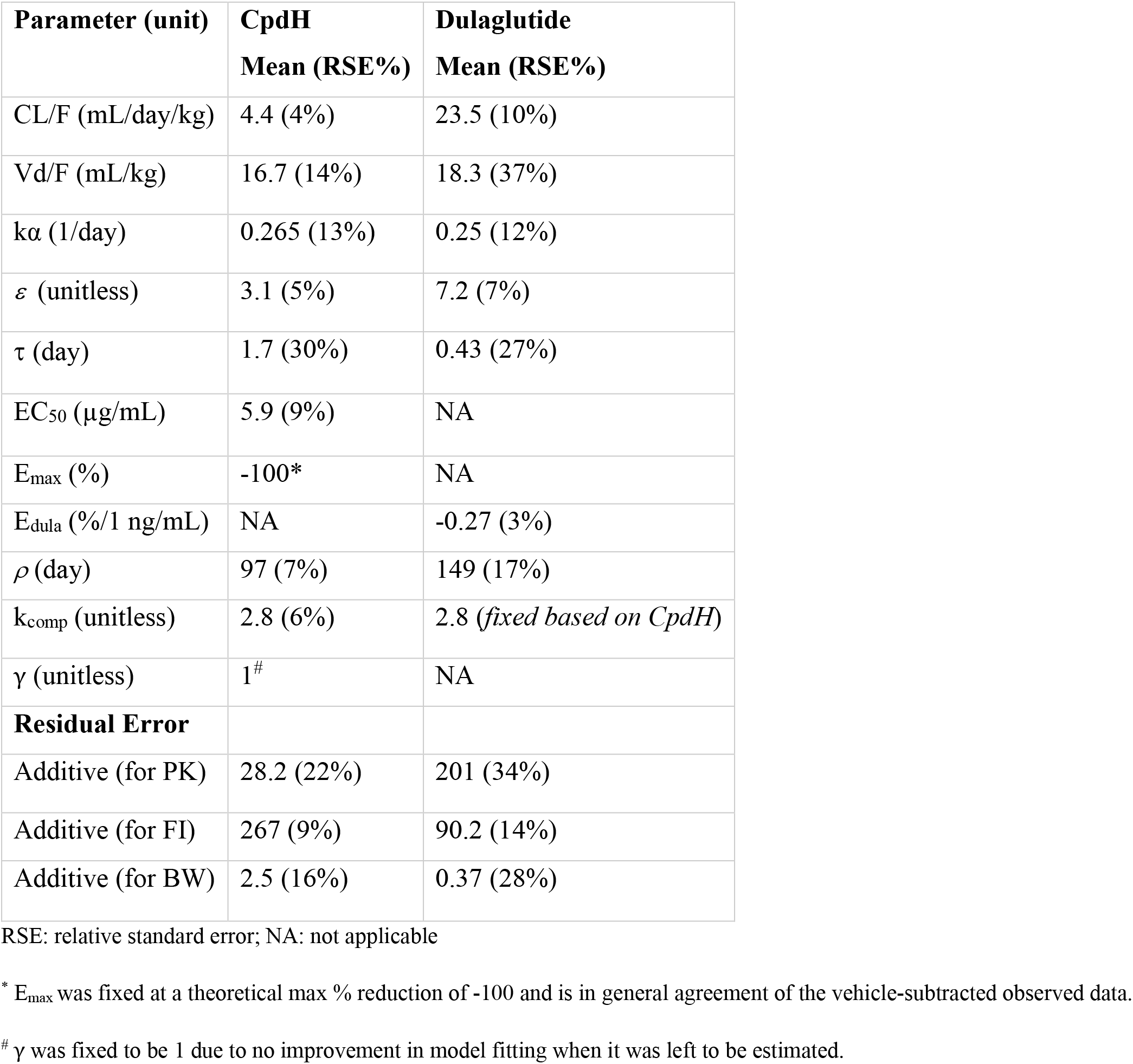
Model-estimated PK/PD parameters for CpdH or dulaglutide in overweight cynomolgus monkeys after repeat dosing

| Parameter (unit) | CpdH<br>Mean (RSE%) | Dulaglutide<br>Mean (RSE%) |
| --- | --- | --- |
| CL/F (mL/day/kg) | 4.4 (4%) | 23.5 (10%) |
| Vd/F (mL/kg) | 16.7 (14%) | 18.3 (37%) |
| $k\alpha$ (1/day) | 0.265 (13%) | 0.25 (12%) |
| $\varepsilon$ (unitless) | 3.1 (5%) | 7.2 (7%) |
| $\tau$ (day) | 1.7 (30%) | 0.43 (27%) |
| EC<sub>50</sub> (µg/mL) | 5.9 (9%) | NA |
| E<sub>max</sub> (%) | -100* | NA |
| E<sub>dula</sub> (%/1 ng/mL) | NA | -0.27 (3%) |
| $\rho$ (day) | 97 (7%) | 149 (17%) |
| k<sub>comp</sub> (unitless) | 2.8 (6%) | 2.8 (*fixed based on CpdH*) |
| $\gamma$ (unitless) | 1<sup>#</sup> | NA |
| **Residual Error** |  |  |
| Additive (for PK) | 28.2 (22%) | 201 (34%) |
| Additive (for FI) | 267 (9%) | 90.2 (14%) |
| Additive (for BW) | 2.5 (16%) | 0.37 (28%) |

RSE: relative standard error; NA: not applicable
\* E<sub>max</sub> was fixed at a theoretical max % reduction of -100 and is in general agreement of the vehicle-subtracted observed data.
<sup>#</sup> $\gamma$ was fixed to be 1 due to no improvement in model fitting when it was left to be estimated.

**Figure 1:**
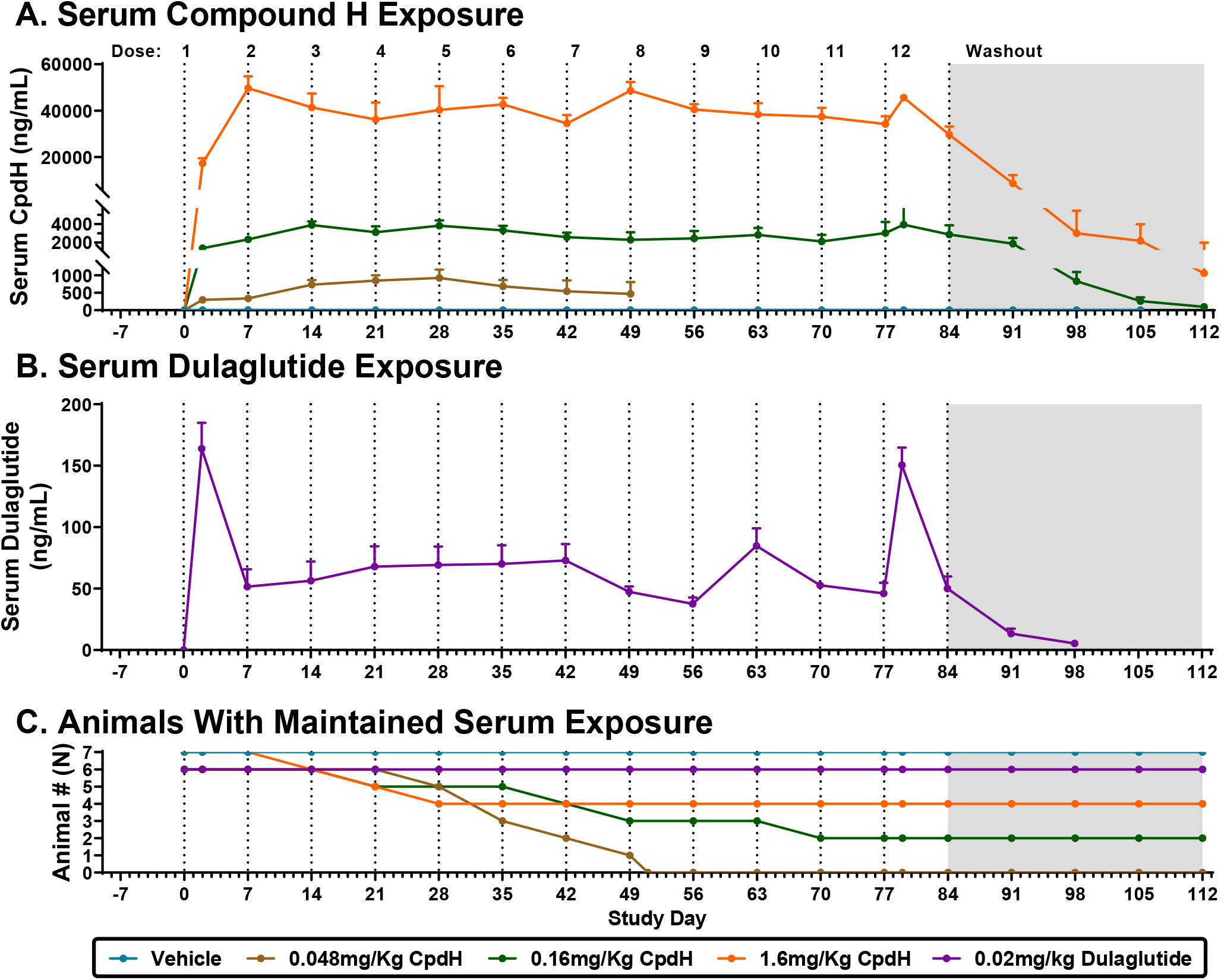
Spontaneously obese cynomolgus monkeys were treated subcutaneously with CpdH (A) or dulaglutide (B) once weekly for 12 weeks, followed by a 4-week washout period. Serum concentration of CpdH and dulaglutide were measured by MSD immunoassay and ELISA, respectively. Administration of an exogenous biologic was anticipated to elicit loss of exposure due to immune activation, which were *a priori* excluded from further analysis according to the data analysis plan; any animal showing loss of >40% measured compound exposure relative to the previous measurement was removed from that point of the study onward. Pharmacodynamic effect of CpdH was rapidly reversed in animals that lost compound exposure (data not shown). The number of animals at each timepoint in panels A and B are reflected in panel C. Data are mean ± SEM.

**Figure 2:**
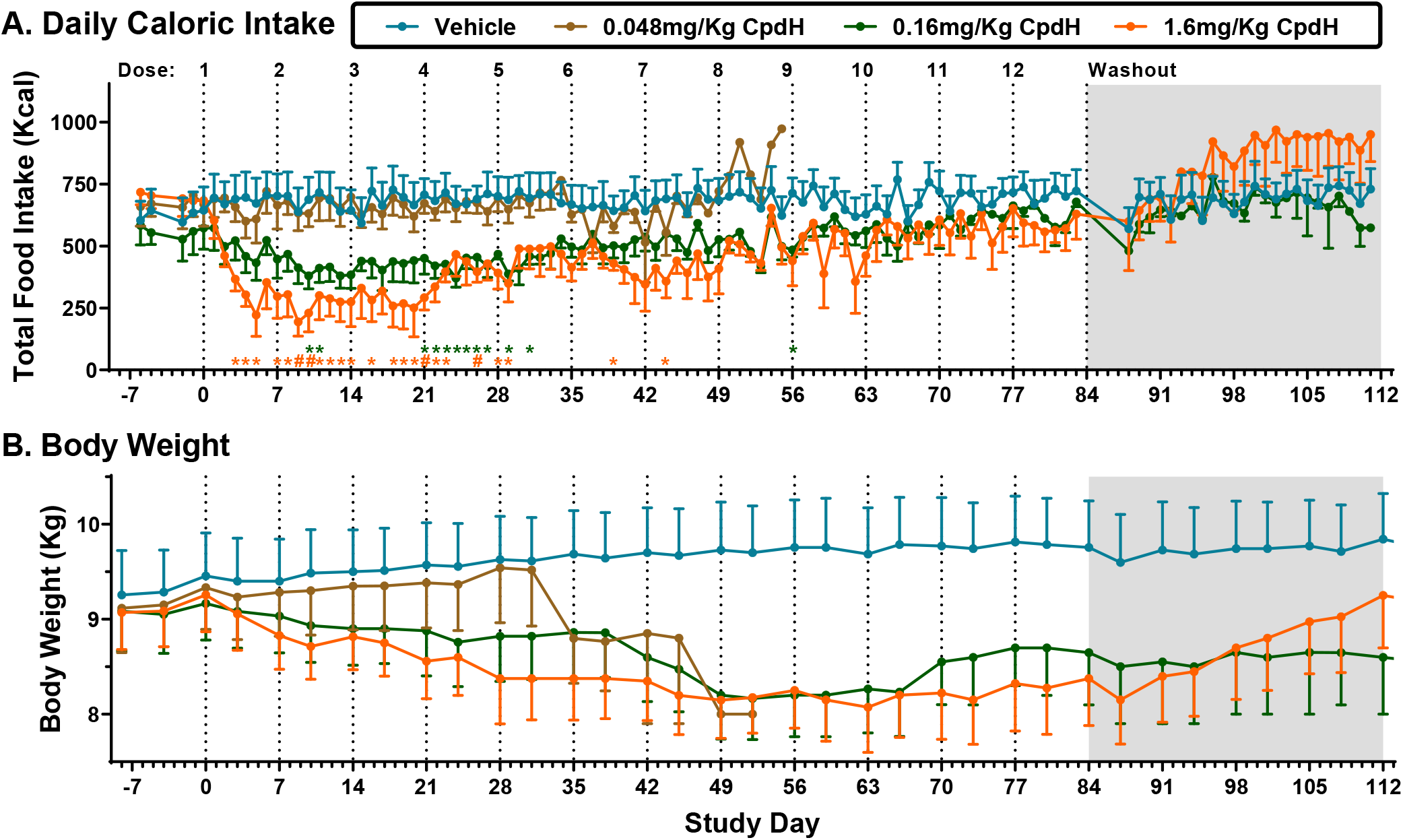
Spontaneously obese cynomolgus monkeys were treated subcutaneously with CpdH once weekly for 12 weeks, followed by a 4-week washout period. Daily caloric intake (A) was measured as the sum of consumed balanced diet provided and the estimated caloric intake derived from the weight and source of daily nutritional enrichment provided. Body weight (B) was recorded twice weekly. The number of animals at each timepoint in panels A and B are reflected in Figure 1C; animals with loss of serum compound exposure were excluded from analysis onward according to the data analysis plan (see Methods section). Data are mean ± SEM; *P<0.05; #P<0.01.

**Figure 3:**
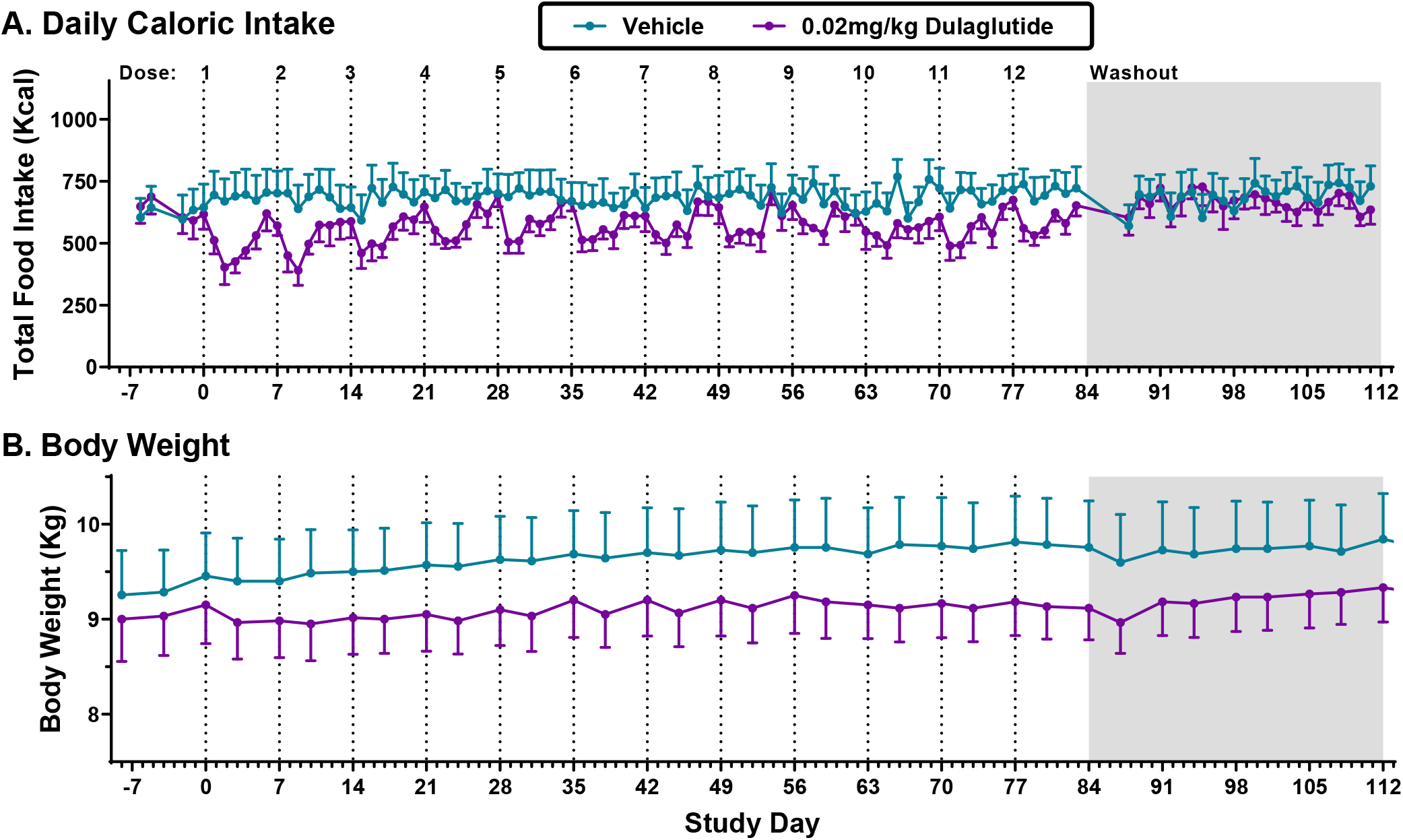
Spontaneously obese cynomolgus monkeys were treated subcutaneously with dulaglutide once weekly for 12 weeks, followed by a 4-week washout period. Daily caloric intake (A) was measured as the sum of consumed balanced diet provided and the estimated caloric intake derived from the weight and source of daily nutritional enrichment provided. Body weight (B) was recorded twice weekly. No loss of dulaglutide exposure was observed during the 12-week treatment phase. Data are mean ± SEM of 7 or 6 animals (vehicle and dulaglutide, respectively).

### Food Intake and Body Weight

Chronic treatment of cynomolgus monkeys with CpdH reduced food intake and body weight relative to vehicle treated animals (**Figure 2**). The 0.048 mg/Kg dose of CpdH was included to model the potentially minimal efficacious dose in human, and did not significantly affect food intake or body weight in the current report (P>0.05 at all timepoints; **Figure 2**). Weekly administration of 0.16 mg/Kg CpdH reduced ad libitum caloric consumption to a nadir of ~24% of baseline after 2 weeks, and continued to suppress food intake for at least 4 weeks (**Figure 2A**). 1.6 mg/Kg CpdH inhibited caloric intake by approximately 54% over the first 3 weeks before rebounding to a new equilibrium level of approximately 22% below baseline for the remaining 9 weeks of treatment (**Figure 2A**). The suppression of food intake by CpdH caused a dose-dependent reduction in body weight (**Figure 2B**), which was sustained during chronic treatment at steady state compound exposure levels. Vehicle-adjusted average body weight reduction for latter 9 weeks of treatment period was 7.1±0.2 and 14.3±0.4% for 0.16 and 1.6 mg/Kg CpdH treatments, respectively. Daily clinical observations throughout the study reported temporary “poor appetite” in 6 monkeys (5 in the 1.6 mg/Kg CpdH group, 1 in the vehicle group), consistent with the mechanism of action of CpdH as an anorectic agent; in all cases of poor appetite, enrichment diet was fully consumed. No other compound-related adverse events were observed, such as emesis, signs of nausea or malaise, which have been reported for other agents impacting satiety; no significant changes in blood chemistry parameters in drug-treated animals were observed (data not shown).

Dulaglutide was included in this study to validate the spontaneously obese cynomolgus monkey model using a clinically relevant agent acting at least in part via food intake suppression. Each weekly injection of dulaglutide caused a rapid reduction in food intake that returned towards baseline near the end of the dosing interval (**Figure 3A**). Notably, the greatest reduction in food intake was approximately 33% following the first two doses of GLP-1 agonist, but the effect attenuated over time. When vehicle corrected, chronic dulaglutide caused a 3.8% reduction in body weight over 12 weeks (**Figure 3B**).

### PK/PD Modeling of Compound H and Dulaglutide in Overweight Cynomolgus Monkeys

The developed PK model provided a reasonable fit to the observed mean PK profiles for both CpdH (**Figure 4A**) and dulaglutide (**Figure 5A**) during the dosing period and the washout period after the last dose, with corresponding estimated PK parameters shown in **Table 1**. The subsequent PD modeling for CpdH was able to simultaneously characterize the PD (% change in EI and BW compared to baseline) of CpdH at the three studied dose levels (**Figure 4B and C**). The modeling results support that the weight loss with CpdH was driven by exposure-dependent reductions in EI without any direct drug effect on EE, consistent with the current knowledge of the mechanisms of action of GDF15. The modeling suggests that the partial rebound in food intake over time is due compensatory changes in EI that occur in response to weight loss (through the kcomp parameter) rather than a loss of drug activity over time, as the parameters describing drug effects on food intake remained constant over time, with an estimated EC50 of approximately 5.9 μg/mL, or ≈ 36 nM for the drug effect on EI.

**Figure 4:**
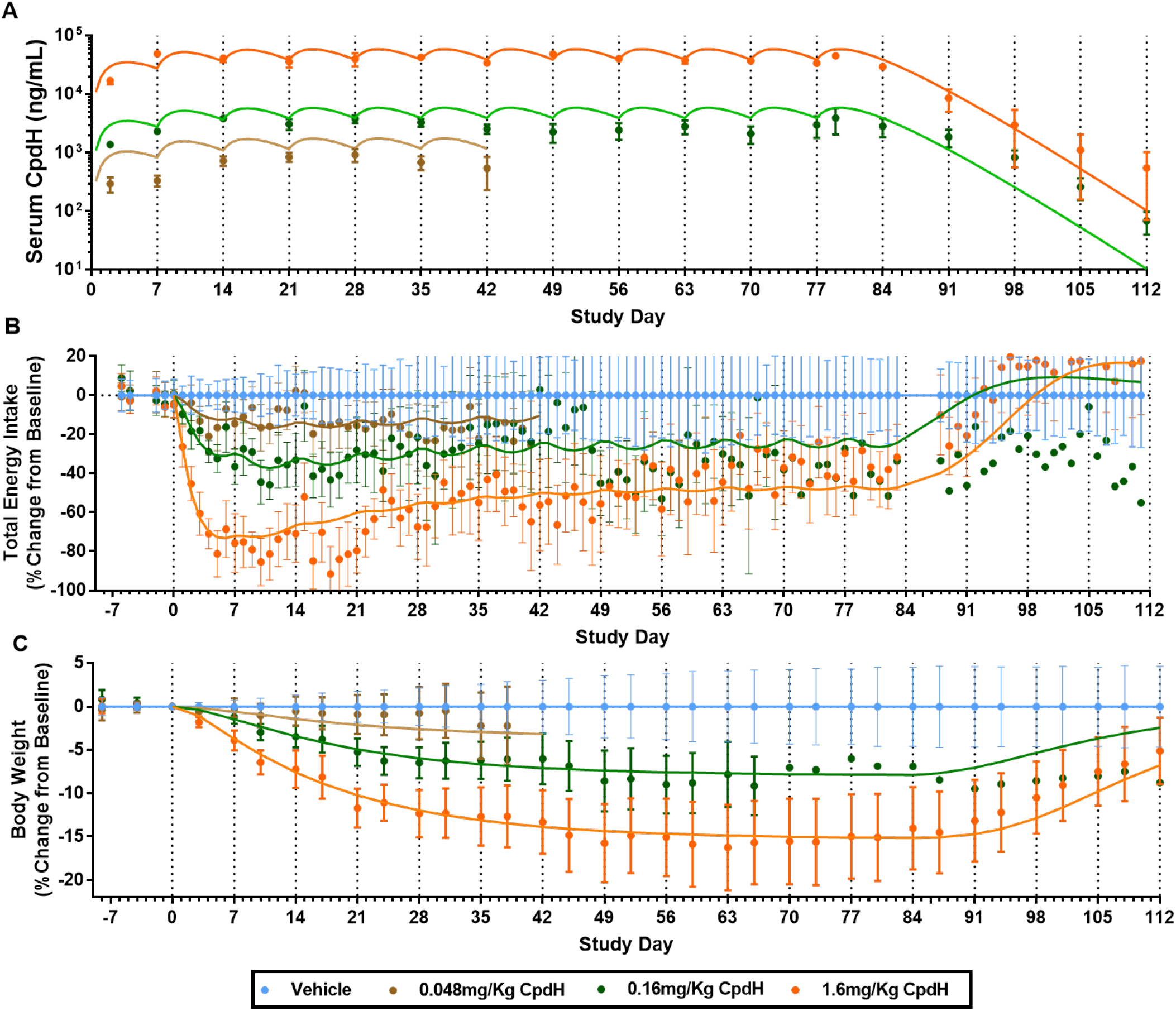
Serum CpdH drug concentrations (A), baseline-normalized, vehicle-subtracted changes in energy intake (B) and body weight (C) in overweight cynomolgus monkeys treated with Compound H. Symbols show the observed mean±SEM and curves show the model fits.

**Figure 5:**
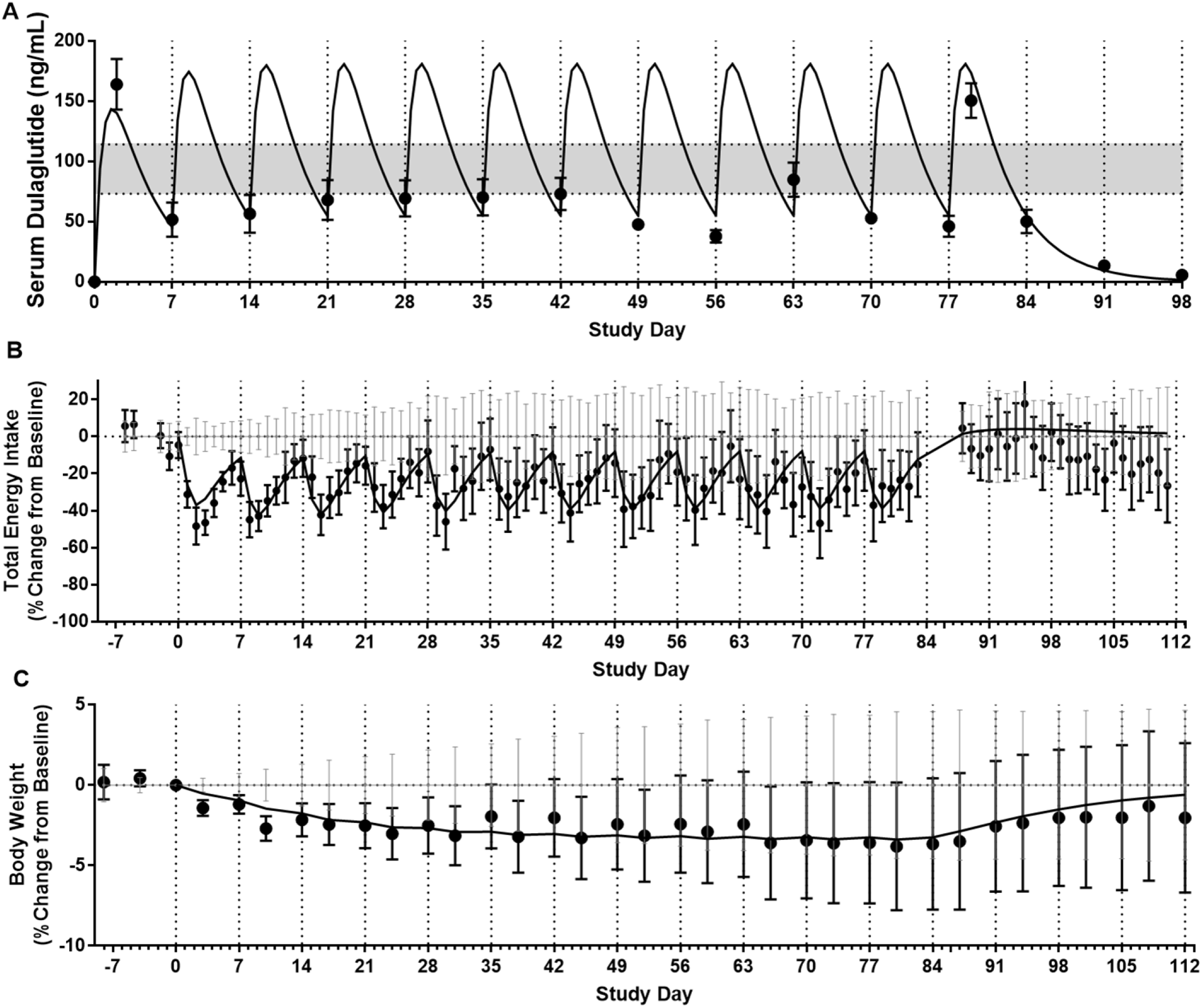
Serum dulaglutide drug concentrations (A), baseline-normalized, vehicle-subtracted changes in energy intake (B) and body weight (C) in overweight cynomolgus monkeys treated with dulaglutide. Symbols show the observed mean±SEM and curves show the model fits. The shaded area in (A) represents the observed mean Cmax and Cmin at steady state with 1.5mg QW SC dosing of dulaglutide in T2DM patients (20).

A similar PD model structure as employed for CpdH, with one adjustment of the E_max_ model to a linear model, was able to simultaneously characterize the PK/EI/BW relationship of dulaglutide in the same study (**Figure 5B and C**). For dulaglutide treatment, the partial rebound in EI within each once-weekly SC dosing interval appears to be primarily explained by drug PK. Model fitting also suggested that the dulaglutide effect on EI is approximately linear within the ≈ 3-fold concentration ranges during each dosing interval, with each 100 ng/mL increase in drug concentration associated with ≈ 27% EI reduction.

The model-estimated PK/PD parameters for both CpdH and dulaglutide are listed in **Table 1**. The parameter ρ linking changes in energy balance to changes in body weight was estimated to be within 1.5-fold for both CpdH and dulaglutide (97 vs. 149, respectively) whereas the parameter *ε* relating changes in EE to changes in BW was estimated to be slightly higher with dulaglutide than with CpdH (7.2 vs. 3.1, respectively). The estimated delay time (*τ*) between systemic drug concentration and each drug’s effect on EI was shorter for dulaglutide than CpdH (0.4 vs. 1.7 days, respectively). All PD parameters for both compounds were estimated with % residual standard error (RSE) ≤ 30%.

### Simulation of dulaglutide’s weight loss effect in humans based on PK and PD parameters estimated in overweight cynomolgus monkeys, and human simulation for Compound H

To assess the translatability of the PK/PD approach for predicting human weight loss based on data from cynomolgus monkeys, we utilized the dulaglutide data in this study along with clinical data describing the human PK and long-term weight loss seen with dulaglutide. Using the modeled human PK (Figure 6A; (20, 21)) together with the PK/PD relationship established in monkeys, predicted human EI reductions with dulaglutide within the range of what was reported in humans with other GLP-1 agonists (22–24) (Figure 6B; no known reports of human food intake reductions with dulaglutide are available). The model predicted long-term reductions in BW were generally in line with reported values for once-weekly SC dose of 1.5mg dulaglutide (≈2.5% vs. 2% for predicted vs. observed, respectively, (20, 25)). Note that the predicted time to reach a plateau in weight loss (≈ 8 weeks; Figure 6C) was shorter than what was typically observed in humans (26).

**Figure 6:**
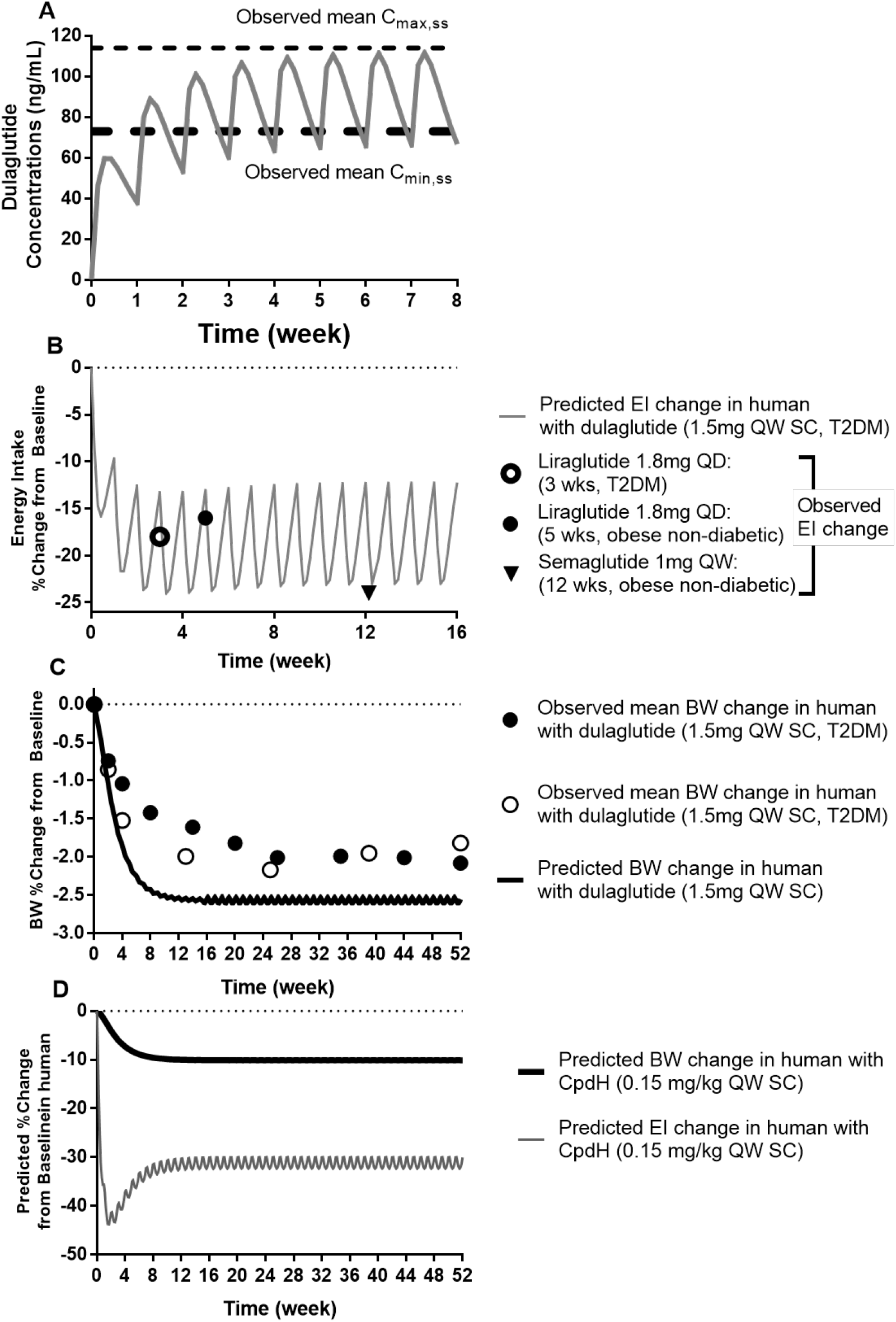
Simulated dulaglutide PK (A) and its impact on EI (B) and BW (C, open circle (35) and solid circle (25)) in human using the dulaglutide PD parameters estimated from overweight cynomolgus monkeys; simulated %change from baseline for EI and BW with CpdH treatment in human (D) based on allometrically scaled PK and the CpdH PD parameters estimated from overweight cyno.

The translational modeling of dulaglutide verified the utility of the obese cyno experimental model and the developed mathematical approach. We used similar approaches for predicting human efficacious dose (see **Methods**) and the long-term weight loss potential of Compound H. **Figure 6D** shows the predicted mean EI and BW profile (% change from baseline) with 0.15 mg/Kg QW SC administration of CpdH over 52 weeks. A steady state of ≈30% EI reduction was predicted which yielded ≈10% BW loss based on the cyno PK/PD relationship.

## Discussion

Increasing prevalence of obesity worldwide has prompted identification and development of novel weight loss therapies. We and others have sought to characterize a circulating half-life extended GDF15 analog in pre-clinical models to validate the central GDF15 receptor GFRAL as a potential drug target. Acute single dose treatment of obese cynomolgus monkeys with HSA-GDF15 caused a dose and time-dependent reduction in food intake with concomitant weight loss (14); an Fc-conjugated GDF15 suitable for once weekly injection in humans reduced body weight over 6 weeks of treatment in a similar obese monkey model (15). Here we demonstrate sustained efficacy of an HSA-GDF15 chimeric analog, CpdH in naïve spontaneously obese NHP. Over a 12-week period, CpdH produced a dose-dependent effect to reduce ad libitum food intake, resulting in sustained body weight loss (**Figure 2**); the comparator dulaglutide similarly produced a significant reduction in body weight over the course of treatment (**Figure 3**). Combined with serum compound exposure data (**Figure 1**), the current study aimed to generate an allometric translational model from monkey to man, to guide human clinical trial study design and dose selection for obesity drugs such as CpdH and other novel weight loss agents.

The PK of CpdH and its impact on the dynamics of EI and BW reduction were well described using a single set of parameters across all three dose levels of CpdH in overweight cynomolgus monkeys and using a separate set of parameters for dulaglutide. The PK/PD modeling approach developed for our studies was built on previous modeling work by Hall and colleagues in humans (27, 28) and related work by Thompson and colleagues in cynomolgus monkeys treated with a Fibroblast Growth Factor 21 (FGF21) analog (29). The work here provides an integrated PK/PD model that accounts for both acute, exposure-dependent effects of treatments to reduce EI (similar to what was done in (29) for 4 weeks of treatment) and also models the attenuation of the pharmacodynamic effect on EI seen with longer treatment. As shown in Hall et al (28), the attenuated effects on EI with sustained treatment and weight loss is assumed to arise from compensatory signals proportional to weight loss rather than a time-dependent loss in the potency or maximal effect of the drug effect on EI. An alternative modeling approach would have been to assume there is tachyphylaxis associated with the drugs’ activity so that the Emax and/or EC50 vary over time. While this alternative hypothesis for the attenuated effect on EI reductions over time cannot be ruled out based on these data, the modeling approach used is consistent with known physiology of body weight regulation and enabled the time profiles for both EI and BW to be described using only a few time-invariant parameters.

As non-human primates are often used to test experimental treatments for therapeutic efficacy, it is of interest to compare the model-estimated parameters relating changes in EI to changes in BW to the corresponding model parameters in humans. In humans undergoing weight loss, EE decreases by approximately 25 Kcal/day per Kg of weight loss (30,31) and increases in appetite were estimated to be ~100 Kcal/day per Kg of weight loss from studies with SGLT2 inhibitor treatment (29) and similar estimate was made using data from several appetite suppressing medications (28). To convert these absolute changes into percentage terms for comparison with the monkey values estimated in this study, baseline EE ~ 2600 kcal/day and baseline BW~100 kg were assumed for a representative obese human; using these values, the 25 Kcal/Kg/day reduction in EE translates into ε_human_ ~ 1 and *k_comp,human_* ~ 4 (i.e., each 1% reduction in BW in humans translates into a 1% reduction in EE and a 4% increase in appetite). For comparison, the estimated ε_cyno_ from the current studies was ~3.1-7.2 with CpdH and dulaglutide, respectively, and due to the relatively small amount of weight loss seen with dulaglutide treatment, the value of ε_cyno_~3 from the CpdH studies is likely more reliable. The estimated value of ε_cyno_~3 is also consistent with results from a previous study reporting reductions in EE in monkeys of 162 Kcal/day per Kg of weight loss (29), which converts to 3.2% reduction in EE per 1% reduction in BW (i.e., ε_cyno_=3.2) using estimated baseline EE ~ 500 Kcal/day and baseline BW~ 10 kg). Thus, weight loss appears to induce approximately 3-fold greater relative reductions in EE in cynomologus monkeys than in humans (3% reduction in EE for each 1% weight loss in cyno vs. 1% reduction in EE in humans). In contrast, weight loss appears to induce generally similar (but modestly lower) increases in appetite in cynomologus monkeys compared to humans based on the estimated *k_comp,cyno_* = 2.8 compared with the estimated *k_comp,human_* ~ 4 (i.e., cynomologus monkeys increase appetite by 2.8% for each 1% decrease in BW vs. 4% increase in humans).

Together, these data suggest that cynomolgus monkeys have a considerably greater reduction in EE in response to weight loss compared with humans, whereas both humans and monkeys have a similar magnitude of compensatory increases in EI in response to weight loss. When the parameters are expressed in percentage terms as done here, the steady-state reduction in BW with sustained treatment will be approximately equal to the initial percentage reductions in EI divided by (ε + *k_comp_*). Thus, if a drug produces a similar initial reduction (e.g. 40%) in EI in both monkeys and humans, the expected steady-state weight loss will be ≈20% greater in humans than in monkeys (e.g., ~8% vs. ~7%, respectively) as ε + *k_comp_* is ~5 in humans vs. ~6 in monkeys. In addition, the time constant associated with the weight loss profile is equal to ρ/(ε + *k_comp_*) and is estimated to be approximately 3 times longer in humans compared to monkeys, consistent with data showing that weight loss in humans typically plateaus somewhere between 16-50 weeks (see e.g. (26)) whereas weight loss in these monkeys plateaued at around 8 weeks.

Finally, using the PK/EI/BW relationship estimated for dulaglutide in overweight cynomolgus monkeys, the simulated steady-state mean BW loss slightly over-predicted the observed human BW data after once-weekly 1.5mg dulaglutide treatment in T2DM patients (~2.5% vs. ~2% for simulated vs. observed mean %BW loss, **Figure 6C**). We therefore took a similar approach for predicting the human PK/PD of CpdH. Although the magnitude and duration of long-term weight loss in humans treated with CpdH is expected to depend on both the magnitude of the initial reductions in energy intake and the extent to which these become attenuated over time, which cannot be readily predicted based on the 12-week NHP study, our simulation (eg, 0.15 mg/Kg QW SC dosing resulted in >30% yearly average EI reduction and ≈10% BW reduction) provided strong support that additional human testing of CpdH is warranted. In studies where energy intake reduction is the sole or main mechanism for weight loss, it has been demonstrated that a mean decrease in energy intake of ≈15% over 52 weeks will result in a 10% BW change (26, 27, 32). If the PK/EI relationship were translatable between cyno and human for CpdH, there is great potential for CpdH to reach >10% weight loss in human at clinically feasible SC doses through a novel mechanism of action.

## Materials and Methods

### Animals

A cohort of 40 spontaneously overweight, biologic agent naïve Adult *Macaca fascicularis* (cynomolgus) male monkeys (8-20 years old, 7.7-11.7Kg) were recruited by Crown Biosciences (Taicang, China). Baseline daily food intake, body weight, DEXA body composition, glycemia, and clinical observations collected over 3 weeks were used to select 32 animals to be used on study; animals were distributed to 5 groups using IRINI software (33) on the basis of average baseline food intake and body weight, body fat, and age. Animals were singly housed in a controlled environment room (20-23°C; 40-70% humidity) under a 12/12h photoperiod. Animals were provided standard NHP balanced diet (Beijing Keao Xieli Feed Company, Ltd, Beijing, China; Protein Content ≥ 16.0, Fat Content ≥ 4.0) twice daily and once daily dietary enrichment (apple, banana, carrot, cabbage, pear or cucumber), and food consumption tabulated. Free access to water was provided. Body weight was measured twice weekly. Animals were chair trained for subcutaneous test article administration and intravenous blood sample collection. Animals were given vehicle, Compound H (HSA-GDF15) at 0.048, 0.16, or 1.6mg/Kg, or 0.02mg/kg dulaglutide once weekly. Animal health history was documented and clinical observations for each subject were recorded daily. All procedures were approved by the Institutional Animal Care and Use Committee and the facility was AAALAC accredited. Cohort baseline characteristics are shown in Supplementary Table 1.

### Test Articles

Recombinant HSA-GDF15 (>96% purity; <0.1 EU/mg endotoxin; Compound H; CpdH) was produced by Excellgene (Monthey, Switzerland). Bioactivity in vitro and in rodents in vivo was confirmed according to previously published methods (14). Test article identity and concentration were blinded to the investigator; individual weekly coded dosing strength stocks were prepared in sterile vehicle [8% sucrose, 0.04% polysorbate20, pH6.5]. Pharmaceutical grade dulaglutide (Trulicity; Eli Lilly) was included as a translational control.

### Serum Bioanalysis

The concentration of HSA-GDF15 in serum samples was determined using an immunoassay method employing a sandwich format with electrochemiluminescent (ECL) detection. Briefly, diluted standards, quality controls, and samples containing the analyte were combined with buffer containing the capture reagent (biotinylated anti-human GDF15 polyclonal antibody, pAb) in a V-bottom Axygen plate and incubated for 60min. Concurrently the Meso Scale Discovery (MSD) Gold Small-spot streptavidin plate was blocked for 60min. The MSD plate was emptied and the analyte/capture reagent mixture was transferred from the Axygen plate to the MSD plate and incubated for 30min. The MSD plate was washed and the detection reagent (ruthenium-labeled anti-HSA monoclonal antibody, mAb) was added and incubated for 30min. The MSD plate was washed and read on an MSD Sector Imager plate reader.

The concentration of dulaglutide in serum samples was determined using an immunoassay method employing a sandwich format with ECL detection. Briefly, MSD Gold Small-spot streptavidin plate was blocked with biotinylated anti-human GLP-1 monoclonal antibody for 60min. Plate was then washed and blocked with 1% BSA. Diluted standards, quality controls, and samples containing the analyte and detection antibody (ruthenium-labeled goat anti-human antibody) were added to the plate, and co-incubated for 60 min. The MSD plate then was washed and read on a MSD Sector Imager plate reader.

### Data Exclusion Criteria

If animals showed loss of compound exposure greater than or equal to a 40% reduction from the previous measurement in the same animal, the data from that point onward was excluded for the remainder of the study.

### Statistical Analysis

All values are given as mean ± SEM. Food intake and body weight changes over time were assessed using a Mixed-effects Model for repeated measures, comparing treatment effect means to the vehicle mean at each timepoint; post-hoc Dunnett’s multiple comparisons test was used to assess significance by adjusted P-value <0.05. Data were analyzed and plotted using GraphPad Prism (v8.4.3).

### PK/PD Model

The PK/PD model for cynomolgus monkeys was developed using the mean PK, energy intake (EI, calculated based on estimated total calorie data) and body weight (BW) data from **Figures 1, 2** and **3**. A one compartment PK model with first-order subcutaneous (SC) absorption (Equation 1) and first-order elimination (Equation 2) was used to describe the PK of CpdH or dulaglutide.

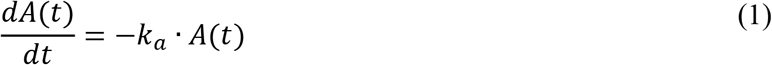

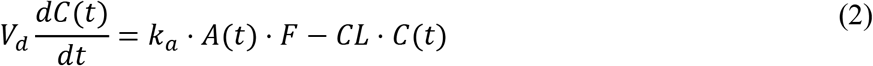

where *A* represents the amount of the drug at the SC injection site, *k*_a_ is the SC absorption rate constant, and *F* is the SC bioavailability. *CL/F* is the apparent clearance, *Vd/F* is the apparent volume of distribution of the drug after SC dosing, and *C(t)* is the drug concentration at time t. Loss of exposure in some animals at later time points was observed for CpdH, but not for dulaglutide, presumably due to development of anti-drug antibodies (ADAs) against CpdH. To compute the mean PK concentrations over time, data from animals that maintained drug exposures were used, by excluding data points when the trough drug exposure was <60% of the previous trough measurement of the same animal. When PK data were below LLOQ, they were treated as missing in the PK/PD modeling. PK parameters were estimated using PK data only and were then fixed during the PD modeling.

The data for both EI and BW were fit using a model that included a concentration-dependent drug effect (*E(t)*) on EI and then a modeled relationship between changes in EI (*δEI*) and changes in BW (*δW*). The relationship between PK and δEI was characterized using an E_max_ (for CpdH) or linear (for dulaglutide) model of drug effect with a first-order time delay between systemic drug concentrations and drug effect with a time constant of *τ* (Equation 3). As described in (27, 28), δW was modeled as being proportional to the difference between EI and EE (Equation 4), and changes in EE (δEE) were assumed to be proportional to δW (Equation 5). Finally, to account for the attenuation of EI reductions over time seen with all pharmacotherapies that reduce EI (likely arising from compensatory signals related to weight loss (28, 30), an additional term proportional to weight loss was included in the equation for EI (Equation 6).

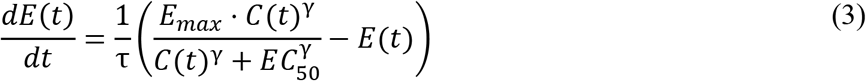

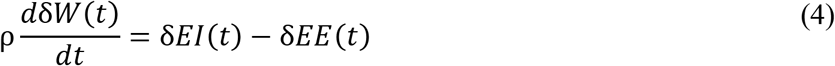

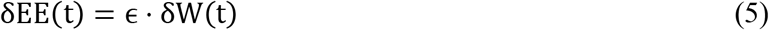

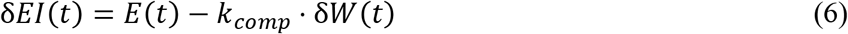

The parameters *ρ*, *ε*, *τ*, *E_max_*, *EC_50_*, and *k_comp_*, were fitted using the measured time profiles for δW(t) and δEI(t). *E_max_* represents the maximum effect of CpdH on reducing EI. *EC_50_* is the drug concentration that gives half-maximal drug effect. *γ*, the Hill coefficient describing the steepness of the concentration–effect relationship was either fixed to 1 or estimated to evaluate its impact on model fitting.

Because only a single dose of dulaglutide was tested, there was not enough information to determine *EC_50_*. Therefore, a linear model linking drug concentration and effect was used for dulaglutide (Equation 3b) and this equation was used in place of Equation 3 in the dulaglutide modeling:

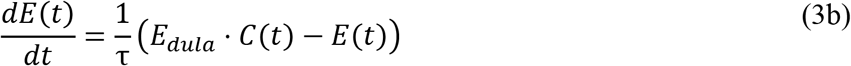

The parameter *k_comp_* describing the weight-loss associated compensatory signal to EI was assumed to be the same for both CpdH and dulaglutide. This parameter could not be estimated well for dulaglutide due to the limited BW change that was seen with the single dose level of dulaglutide studied. Therefore, *k_comp_* was first estimated using CpdH data and was then fixed during the PD model fitting for dulaglutide. Other PD parameters such as *E_dula_*, *ρ*, *τ*, and *ε* were fitted separately for CpdH and dulaglutide.

Fixed-exponent allometric scaling based on body weight (eg, 90 Kg and 9 Kg for human and cyno) was used to predict human PK of CpdH based on overweight cynomolgus monkeys. An exponent value (scaling factor in Equation 7) of 0.75 was used for apparent SC clearance (CL/F; calculated value for human=2.5 mL/day/Kg), and 1 was used for apparent SC volume of distribution (Vd/F; calculated value for human=16.7 mL/Kg). Note that the exponent values for both CL and Vd suitable for human PK prediction are not extensively studied for HSA based drug conjugates, and the translatability of SC absorption between overweight monkeys and overweight humans is unknown. The choice of exponent values was based on other therapeutic antibodies following IV dosing (34). The equation used to perform allometric scaling are provided below in which P represents either apparent CL or Vd:

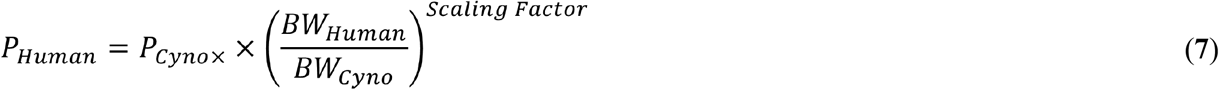

In absence of specific data, the SC bioavailability (F%) and the first-order absorption rate constant Ka (≈0.265 day^-1^) are assumed to be similar between overweight monkeys and overweight humans.

The PK/PD data were analyzed using the nonlinear mixed-effects approach with NONMEM version 7.3.0 (Icon Development Solutions) to obtain population parameter estimates. Because the mean PK and PD data were used, the between-subject variability was fixed to be zero. The residual error structures included additive error for PK and additive error for PD (both δEI and δBW). The first order conditional estimation with interaction (FOCEI) and the ADVAN 13 subroutine were used. Goodness-of-fit plots were used for model evaluation.

## Acknowledgments

The authors would like to express gratitude to Mark Erion, Shamina Rangwala, Vedrana Stojanovic-Susulic, Katharine D’Aquino, Shannon Mullican, James Leonard, Peggy Wong, Ellen Chi, Michael Hunter, Ron Swanson and Holly Kimko for input during experimental design and data analysis. The authors thank the scientists at CrownBio for executing the study as designed.

---

**Supplementary Table 1:**
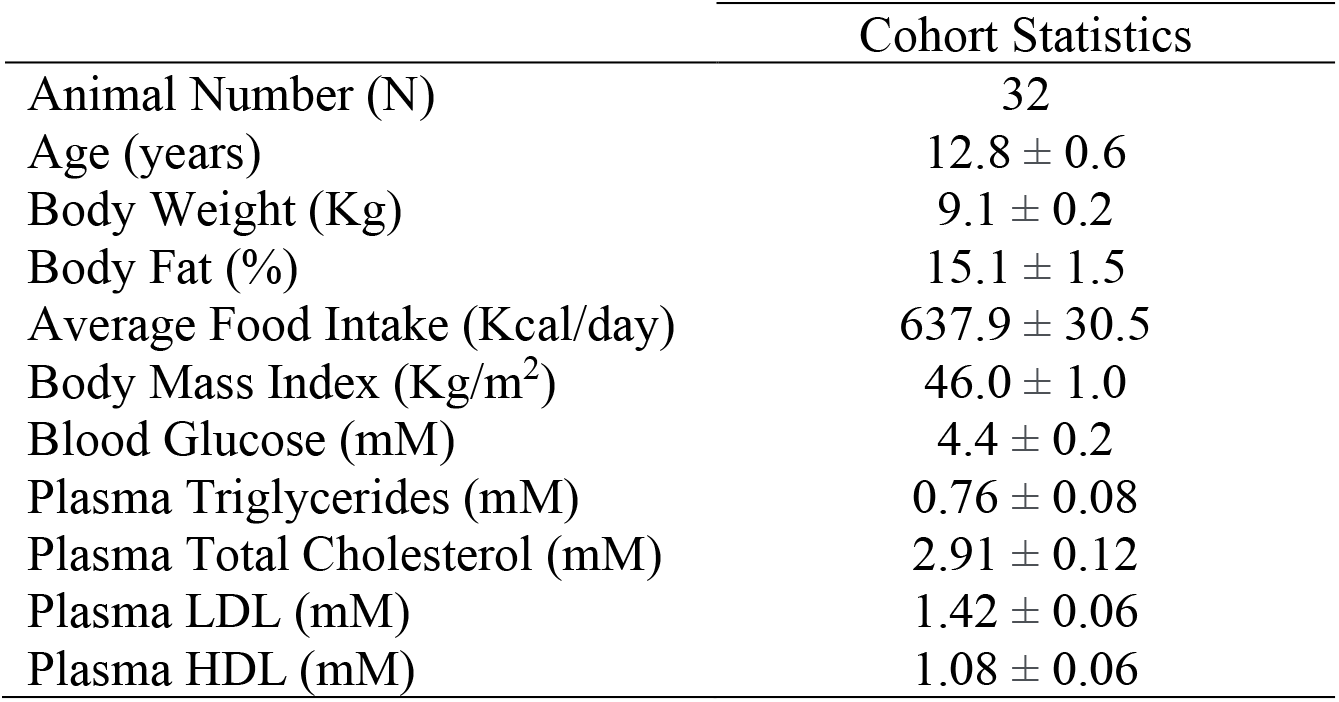
Cynomolgus monkey baseline characteristics. Animals were fasted overnight (16 hours) prior to DEXA body composition analysis and blood chemistry. Average Food Intake represents the average of 5 days prior to compound treatment. Data are mean ± SEM.

**Supplementary Table 2:** Grouped baseline characteristics. Animals were fasted overnight (16 hours) prior to DEXA body composition analysis and blood chemistry. Average Food Intake represents the average of 5 days prior to compound treatment. Data are mean ± SEM.

|  | Vehicle | 0.048mg/Kg<br>CpdH | 0.16mg/Kg<br>CpdH | 1.6mg/Kg<br>CpdH | 0.02mg/Kg<br>dulaglutide |
| --- | --- | --- | --- | --- | --- |
| Animal Number (N) | 7 | 6 | 6 | 7 | 6 |
| Age (years) | 13.6 ± 1.6 | 12.7 ± 1.1 | 12.0 ± 1.1 | 12.7 ± 1.3 | 13.0 ± 1.7 |
| Body Weight (Kg) | 9.3 ± 0.4 | 9.1 ± 0.4 | 9.1 ± 0.4 | 9.0 ± 0.4 | 9.0 ± 0.4 |
| Body Fat (%) | 15.5 ± 3.1 | 15.9 ± 4.0 | 16.1 ± 4.2 | 13.9 ± 2.8 | 14.5 ± 3.8 |
| Body Mass Index (Kg/m<sup>2</sup>) | 44.9 ± 1.9 | 45.4 ± 2.6 | 43.7 ± 1.1 | 49.9 ± 2.3 | 45.8 ± 2.4 |
| Average Food Intake (Kcal/day) | 624.5 ± 82.6 | 672.1 ± 74.0 | 559.5 ± 77.4 | 696.0 ± 50.0 | 629.8 ± 63.3 |
| Blood Glucose (mM) | 4.5 ± 0.4 | 4.4 ± 0.5 | 4.5 ± 0.2 | 4.6 ± 0.4 | 4.1 ± 0.4 |
| Plasma Triglycerides (mM) | 0.58 ± 0.07 | 0.79 ± 0.17 | 0.81 ± 0.25 | 0.94 ± 0.23 | 0.69 ± 0.14 |
| Plasma Total Cholesterol (mM) | 2.81 ± 0.20 | 2.81 ± 0.26 | 2.85 ± 0.36 | 3.02 ± 0.33 | 3.03 ± 0.19 |
| Plasma LDL (mM) | 1.06 ± 0.09 | 1.00 ± 0.13 | 1.08 ± 0.15 | 1.13 ± 0.18 | 1.10 ± 0.09 |
| Plasma HDL (mM) | 1.36 ± 0.12 | 1.39 ± 0.13 | 1.35 ± 0.20 | 1.48 ± 0.14 | 1.54 ± 0.16 |

## Notes

### Competing Interest Statement

All authors are or were employees of Janssen Pharmaceutical Companies of Johnson & Johnson. All authors hold J&J stock.

